# Cyclic stretch regulates epithelial cell migration in a frequency dependent manner via vinculin recruitment to cell-cell contacts

**DOI:** 10.1101/2023.08.19.553915

**Authors:** Liam P. Dow, Stacey Surace, Katrene Morozov, Reagan Kennedy, Beth L. Pruitt

## Abstract

Epithelial cell migration is critical in regulating wound healing and tissue development. The epithelial microenvironment is incredibly dynamic, subjected to mechanical cues including cyclic stretch. While cyclic cell stretching platforms have revealed responses of the epithelium such as cell reorientation and gap formation, few studies have investigated the long-term effects of cyclic stretch on cell migration. We measured the migratory response of the epithelium to a range of physiologically relevant frequencies and stretch. We integrated our experimental approach with high-throughput cell segmentation to discover a relationship between changes in cell morphology and migration as a function of cyclic stretch. Our results indicate that lower stretch frequencies (i.e., 0.1 Hz) arrest epithelial migration, accompanied by cell reorientation and high cell shape solidity. We found that this response is also accompanied by increased recruitment of vinculin to cell-cell contacts, and this recruitment is necessary to arrest cell movements. This work demonstrates a critical role for frequency dependence in epithelial response to mechanical stretch. These results confirm the mechanosensitive nature of vinculin within the adherens junction, but independently reveal a novel mechanism of low frequency stress response in supporting epithelial integrity by arresting cell migration.

## Introduction

Organ function and tissue development rely on interconnected sheets of epithelial cells. In adult tissue, these cells regulate nutrient absorption, filtration, and act as a mechanical barrier against trauma or pathogenic invasion. During tissue development, they fold and constrict, shaping new tissues. Unlike passive cellular materials such as foams, cork, honeycombs, etc., the epithelium consumes and exerts energy as independent units. This property allows epithelial cells to migrate, rearrange, and alter their cellular shape ^1–4^, facilitating their specific functions in dynamic microenvironments ^5^.

Epithelial microenvironments exhibit high dynamism, particularly in terms of mechanical stretch and deformation. For example, lung alveolar epithelial cells experience cyclic stretching due to respiratory rhythms ^6^, while peristaltic contractions of smooth muscle cyclically stretch regions of the intestinal epithelium ^7^. In developing tissues, pulsatile stretches help orchestrate events such as gastrulation and dorsal closure in *Drosophila* ^8,9^. These cyclic mechanical stretches occur across various epithelial microenvironments but differ significantly in frequency and magnitude. Many diseases also disrupt the natural mechanics of epithelia by altering stretch magnitudes and rates, as seen in irritable bowel syndrome ^7^, asthma ^6,10^, and colorectal cancer ^11^.

Several studies have employed *in vitro* mechanical stretching experiments to elucidate the effects of cyclic stretch on epithelial behavior. At higher cyclic stretch amplitudes and frequencies (>20%, >1 Hz), researchers have observed gap formation in epithelia of parental Madin-Darby canine kidney (NBL-2) cells ^12^ and a decrease in wound healing time compared to an unstretched epithelium ^13^. There is also evidence that cyclic stretch mediated wound healing is ECM dependent ^14^. Researchers have also observed epithelial cell shape reorientation in response to uniaxial cyclic stretch ^15–18^ which is believed to influence morphogenesis and wound healing ^19,20^. Studies of isolated NRK (normal rat kidney) epithelial cells (stretched from 5-20% at 0.5Hz) found stretch-induced reorientation depended on microtubule remodeling ^15^, while studies of isolated A549 alveolar epithelial cells (stretched from 5-15% at 0.3 Hz) found this reorientation also required stress fiber remodeling (i.e., F-actin) ^17^. Stretch frequencies have ranged broadly from study to study (0.1 Hz – 2.15 Hz) ^12,14–18^, which may contribute to different findings. Moreover, comparison across these studies is complicated by the use of different strains of epithelial cells which can have different adhesion and proliferation characteristics ^21^. In related experimental studies ^22,23^ and single cell theoretical models ^24^, direct comparisons of the effects of stretch frequency on endothelial or fibroblast stress fiber reorientation do suggest a frequency dependent relationship. However, the role of cyclic stretch frequency in regulating collective epithelial behavior is less clear and we sought to address this gap.

Under in-plane mechanical load, epithelial cells exert forces on each other at their cell-cell contacts. These contacts encompass several junctions, including tight junctions, desmosomes, gap junctions, and adherens junctions ^25–27^. Among these, the adherens junction plays a mechanosensitive role in guiding cell migration ^28^ and regulating cell proliferation ^29,30^, even responding to static stretches ^31–33^. Yet, the role of the adherens junction in regulating responses to cyclic stretch is unknown.

The transmembrane protein E-cadherin serves as the intercellular mechanical linkage in the adherens junction. E-cadherin forms a trimeric complex with cytosolic β-catenin and α-catenin, which binds to F-actin ^25^. Various adapter proteins stabilize the binding of this complex to F-actin under mechanical load. For example, p120-catenin localizes to E-cadherin to block clathrin-mediated endocytosis of E-cadherin and maintain stability of the complex ^34^. Another protein, vinculin, reinforces the α-catenin/F-actin interaction under mechanical load ^35–37^ to fortify the adherens junction ^38^. Despite the known mechanosensitive recruitment of vinculin to the adherens junction, it is unclear how it is recruited with cyclic stretch and how that recruitment affects epithelial shape change and migration.

We sought to learn how different frequencies of uniaxial cyclic stretch impacted epithelial behavior and the potential role of vinculin in mediating this behavior. We subjected high density, confluent Madin-Darby canine kidney (MDCK type II) epithelial monolayers to a range of physiological cyclic stretch frequencies (0.1 Hz, 0.5 Hz, and 1 Hz) at a physiological stretch magnitude of ∼10% ^39,40^. Observing the effect of cyclic stretch on the epithelium over time is challenging because stretch devices must be amenable to long-term live cell imaging; to overcome this challenge, we integrated a programmable pneumatic cell stretching device ^41^ with a microfluidic perfusion system to observe cell migration after cyclic stretch.

## Results

### Cells adjust morphology in response to cyclic stretch

To investigate the long-term dynamic response of the epithelium to cyclic stretch, we utilized a pneumatic cell stretcher device previously detailed ^41^ (**Fig. 1a**). First, we created a polydimethylsiloxane (PDMS) device from a 3-D printed mold and adhered a thin PDMS bottom membrane, forming 2 main channels. The center channel, flanked by inlet and outlet ports, functioned as the substrate for the epithelium. The outer air-filled channel, encircling the center channel, contracted when vacuum was applied. This contraction of the outer channel led to the uniaxial stretching of the membrane of the central channel and the adherent epithelium. By applying a programmable input pressure of 60 kPa, cells were uniaxially stretched ^29,41^, resulting in ε_yy_ = 10% ± 2% as measured by cell membrane elongation (**Supplementary Fig. 1a**). Since this small level of stretch is difficult to observe, stretched cells were labeled with Hoechst to visualize nuclei displacement (**Fig. 1b**). To examine the impact of 10% stretch on cell morphology, we performed a high-throughput automated segmentation of the epithelium before and after stretch (**Supplementary Fig. 1b-d**). This method enabled the analysis of numerous cells, offering robust statistical power to identify variations in cell morphology such as average cell area and perimeter.

**Figure 1.**
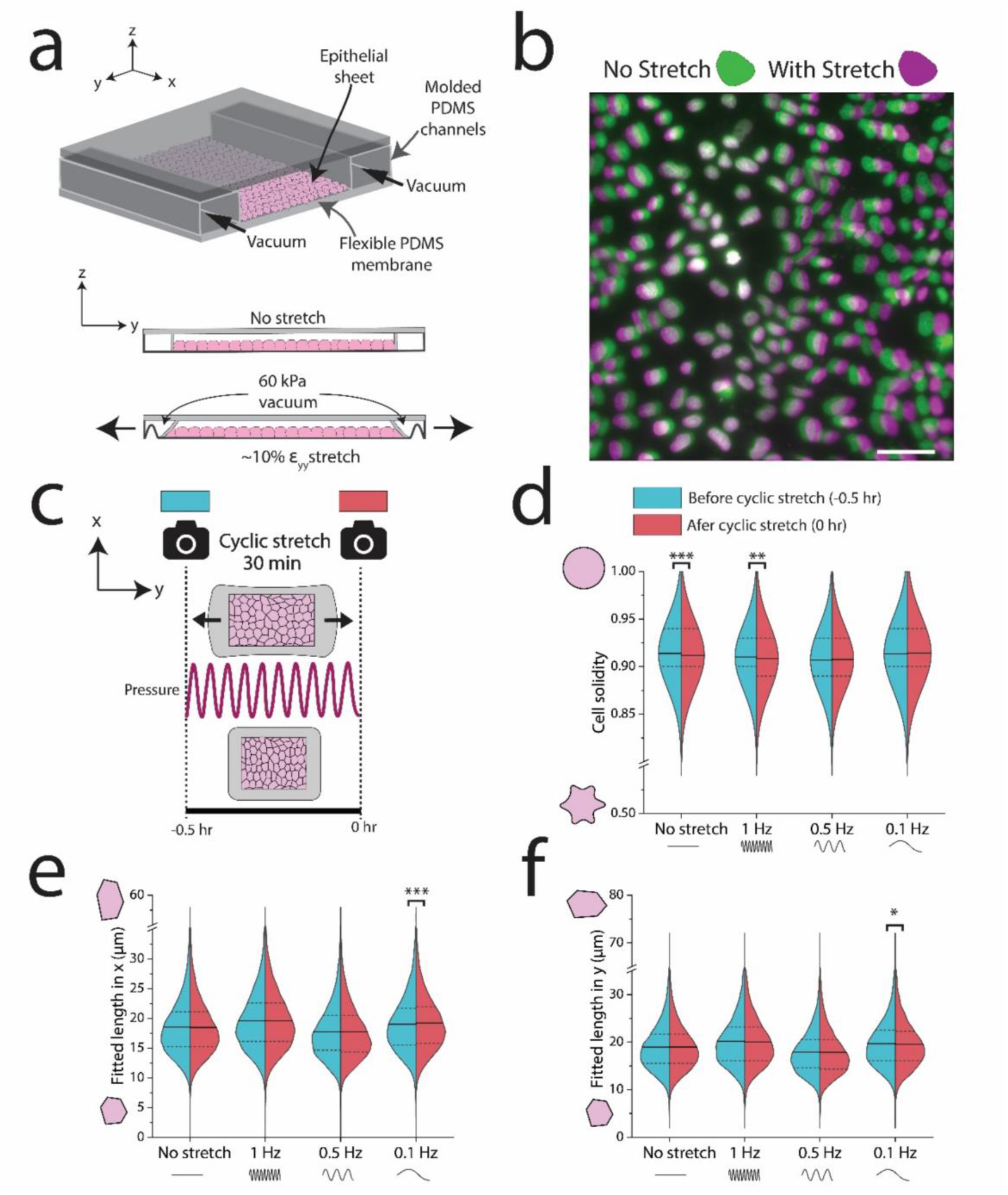
Demonstration of uniaxial stretch device and segmentation of epithelial cells after cyclic stretch. (a) Schematic of PDMS pneumatic cell stretching device. Vacuum in the outer chambers contracts the epithelium ∼10% uniaxially. (b) Nuclei were labeled with Hoechst and imaged before and after application of a 60 kPa vacuum to generate a 10% uniaxial stretch. Green nuclei indicate cells without stretch (0 kPa) and magenta denotes displaced nuclei in stretched cells (60 kPa). (c) Schematic of experiment for applying cyclic stretch to an epithelial sheet, where cells were imaged immediately before and after application of a sinusoidal 10% stretch at different frequencies for 30 minutes. (d) At lower frequencies, cells retained their solidity. (e and f) At 0.1 Hz, cells elongated perpendicular to stretch and contracted in the direction of stretch. n>10,000 cells per half of each violin plot. Scale bar in (b) is 50 µm.

After validating that a 10% static stretch changed cell morphology, we subjected the epithelium to 1 Hz, 0.5 Hz, or 0.1 Hz cyclic stretch of 10% for 30 minutes (**Fig. 1c**). We imaged the epithelium before and after cyclic stretch, with the membrane fully relaxed (time = -0.5 hours and 0 hours, respectively). We then performed a large-scale segmentation of all images, which was made possible by a stably transfected cell line with a GFP E-cadherin fluorophore ^42^. We extracted several shape descriptors from these segmented images, including cell solidity. Solidity is defined as 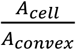, where A cell is the cell area and A_convex_ is the convex area. Higher solidity indicates rounder cells, while lower solidity implies more protrusions ^43^. This metric has been used to assess cell deformability and is associated with age-related macular degeneration of retinal pigment epithelial cells ^44,45^. Interestingly, we found that solidity was highly frequency dependent (**Fig. 1d**). Higher frequency (1 Hz) stretch or no-stretch led to decreased average cell solidity. In contrast, lower frequencies (0.5 Hz and 0.1 Hz) did not elicit changes in average cell solidity, with a subtle increase (though not statistically significant). Cell morphological parameters including circularity and shape index 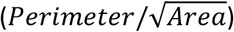 were also disrupted under higher frequency 1 Hz cyclic stretch, but not 0.1 Hz (**Supplementary Fig. S2**).

At the lower frequency of 0.1 Hz, we also observed a reorientation of epithelial cells (**Fig. 1e-g**). Cells reduced their length in the direction of uniaxial stretch (y) and extended their length perpendicular to stretch (x). This shape change corroborates previous studies of how cyclic stretch modulates cell shape ^15,18,46^. Furthermore, we did not observe cell reorientation at 0.5 Hz or 1.0 Hz. These investigations extended the analysis of morphological shifts by linking reorientation to the stabilization of cell solidity. These morphological alterations suggested a role for intercellular signaling at cell-cell contacts.

### Low frequency stretch arrests epithelial migration

After cyclic stretching the epithelium for 30 minutes (time = 0 hours), we observed how the epithelium changed its collective migration over the next 6 hours (**Fig. 2a**). Comprehensive studies of how cyclic stretch governs the collective epithelium are limited, partly due to the challenges inherent in such investigations. Conducting live cell imaging for prolonged periods on a cell stretcher platform is demanding. The bottom membrane must be both transparent and thin enough for imaging, and many cell stretcher setups are not fully enclosed. Open systems are susceptible to media evaporation within hours and cannot be continuously imaged on a microscope. To counteract this challenge, we integrated a microfluidic based perfusion system within our device to prevent evaporation (**Fig. 2b**). A syringe pump sustained a slow media flow rate of 300 µL/hr, resulting in minimal shear stress of approximately 1×10^−6^ dyne/cm^2^ on the apical epithelium (**Fig. S3**). Furthermore, we eliminated out of plane drift during imaging by adhering the device to a thin glass coverslip.

**Figure 2.**
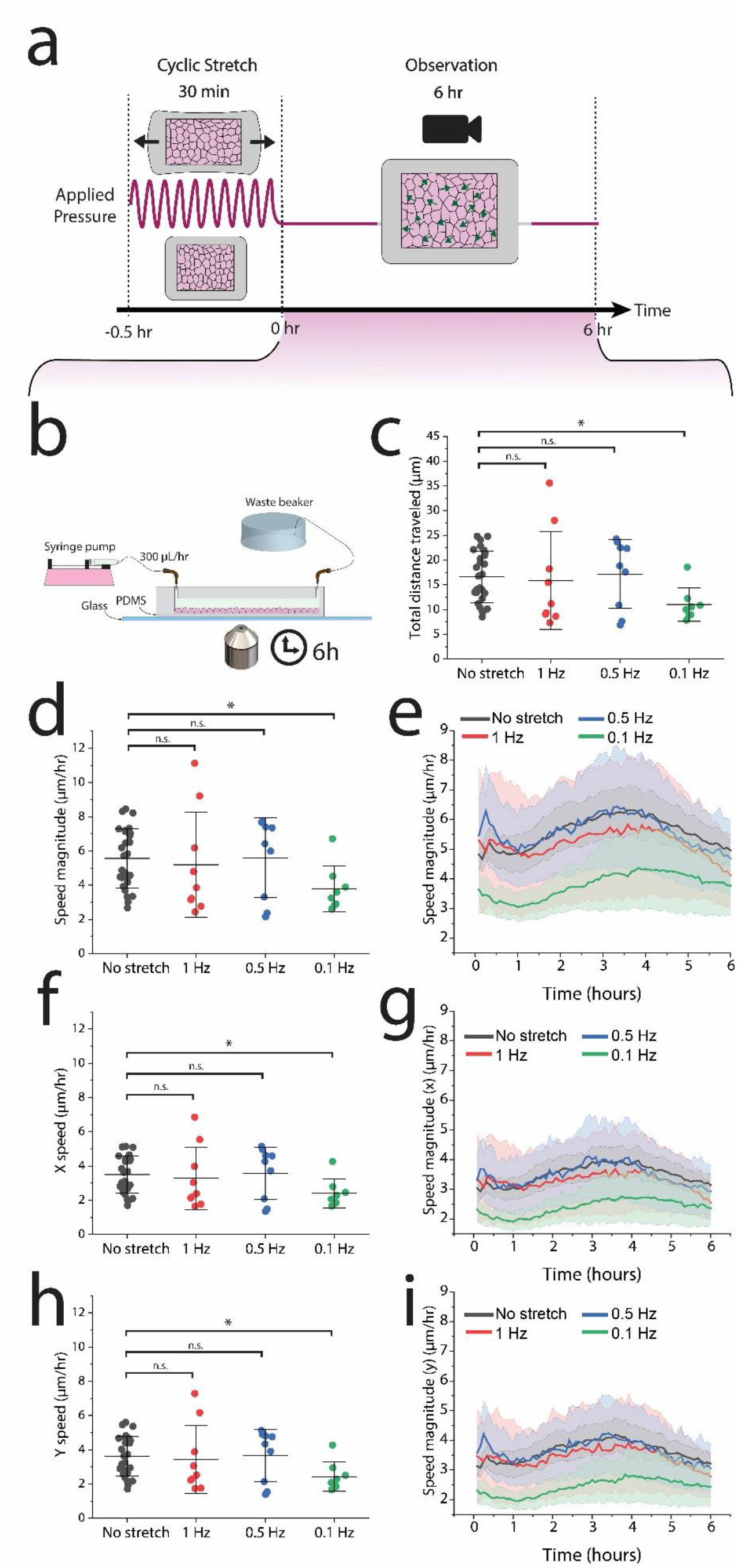
Low frequency cyclic stretch reduces epithelial migration in a directionally independent manner. (a) PIV analysis was conducted in the observation period, the 6 hours following 30 minutes of cyclic stretch. (b) After cyclic stretch, the device was integrated with a perfusion system and imaged on a glass slide to remove out-of-plane drift. (c) Cells traveled the shortest distance in response to 0.1 Hz, with no significant differences among the higher frequencies. (d) The average cell speed diminished significantly in response to a 0.1 Hz frequency. (e) The reduced migratory effect predominantly lasted ∼2 hours. (f-i) Both the x and y speeds are reduced in response to 0.1 Hz uniaxial stretch. No-stretch control: n=27 across 9 independent experiments, 1 Hz: n=9 across 3 independent experiments, 0.5 Hz n=9 across 3 independent experiments, 0.1 Hz: n=8 across 3 independent experiments.

We utilized particle image velocimetry (PIV) to track displacement of cells during this 6-hour period. This method facilitated the calculation of migration parameters including average speed, total distance traveled, and specific x and y movements.

Average speeds for the no-stretch control, 1 Hz, and 0.5 Hz conditions were approximately 5-6 µm/hr. Notably, the slowest frequency condition (0.1 Hz) displayed a significant reduction in average speed to around 3.8 µm/hr (**Fig. 2c**). The total distance traveled aligned closely with the average speed results, with 0.1-Hz stretched cells covering nearly half the distance of cells in the other conditions (**Fig. 2d**).

To discern whether the migratory responses occurred over different timeframes, potentially masking variations in speed averages, we plotted migration speed against time (**Fig. 2e**). Notably, the speed was most significantly disrupted immediately following a 0.1 Hz cyclic stretch. After 6 hours the speed equalized across all frequencies, resembling the no-stretch control. Surprisingly, the migratory response of cells in the 0.5 Hz and 1 Hz conditions closely matched the migration timescales of the control condition. In all cases, we observed a sigmoidal response in speed fluctuation over time, with initial deceleration, brief acceleration, and subsequent deceleration towards the end of the observation period.

Given the uniaxial nature of the applied stretch (ε_yy_), we also explored the possibility of direction-specific migration dysregulation. However, migration was slowed *both* the direction of stretch (y) and perpendicular to stretch (x) (**Fig. 2 f-i**).

### Adherens junction regulates arrest of cell movements

After establishing that slow frequencies disrupted collective epithelial migration speed, we investigated the role of an intact adherens junction in regulating this migration reduction. Therefore, we utilized MDCK cells expressing a mutant E-cadherin protein (T151 cells, made in an MDCK GII background), lacking the extracellular domain of E-cadherin, thus inhibiting adherens junction interaction ^47^. The T151 MDCK cells have a doxycycline (DOX) repressible promoter ^47^; they maintain a wild-type (WT) E-cadherin ectodomain when cultured with DOX. Removal of DOX from the media causes the E-cadherin to mutate, leading to the absence of the outer ectodomain ^47^. The presence of the mutated E-cadherin inhibits mechanical linkages at the adherens junction (**Fig. 3a**), though cells are still able to attach via desmosomes and tight junctions.

**Figure 3.**
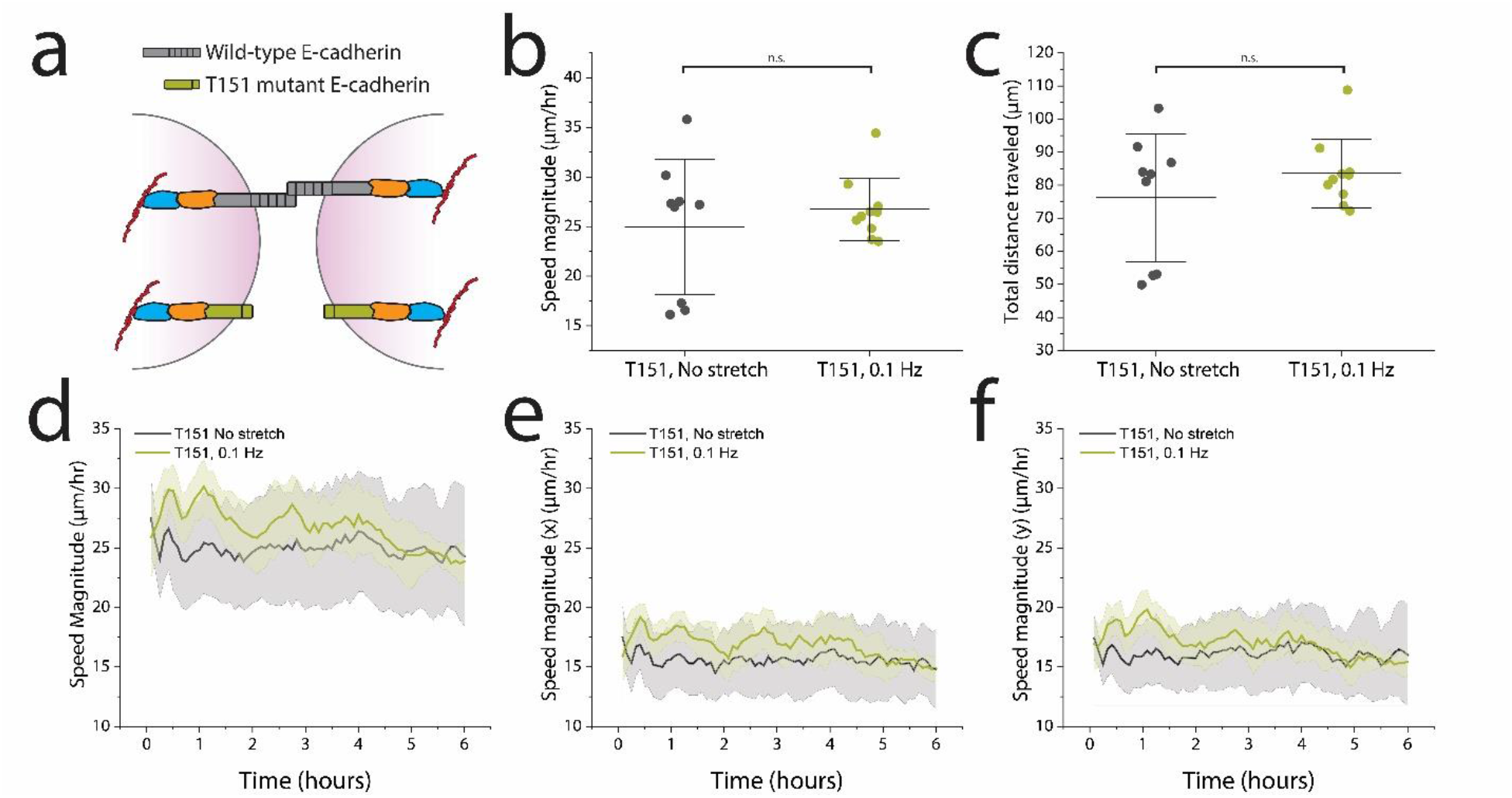
The extracellular domain of E-cadherin is necessary for reducing migration after 0.1 Hz stretching. (a) The extracellular domains of E-cadherin are truncated on the T151 MDCK cells, preventing formation of the adherens junction while allowing formation of other cell-cell adhesions. (b,c) T151 MDCK cells showed no significant differences in collective average speed magnitude nor total distance traveled over the course of 6 hours following 0.1 Hz cyclic stretch (green). (d) Migration speed after cyclic stretch did not differ significantly from the no-stretch control. (e,f) The migratory response showed no directional dependence. No-stretch control: n=9 across 9 independent experiments, 0.1 Hz: n=10 across 4 independent experiments.

We noted distinctive characteristics of the T151 epithelium compared to WT MDCK epithelium. First, in our hands the epithelium did not grow as dense. Second, small holes can form in the epithelium over time, a phenomenon which has been previously reported ^47^.

Overall, we found that T151 cells were more migratory and maintained a faster average migration speed of approximately 25 µm/hr. Despite the increase in migration speed compared to the wild-type MDCK cells, we observed no differences in migration between T151 cells subjected to 0.1 Hz cyclic stretch and those without cyclic stretch (**Fig. 3b, c**). Furthermore, migration alterations over time were not significant (**Fig. 3d-f**). Immunohistochemistry (IHC) staining of the devices confirmed the lack of an adherens junction complex, as evidenced by minimal p120-catenin expression at cell-cell contacts (**Fig. S4**). Together, these results indicate that an intact adherens junction is necessary for reduced migration following 0.1 Hz cyclic stretch.

### Cell-cell contacts recruit vinculin in response to low frequency stretch

After establishing the stretch dependent migratory response of the adherens junction, we looked at the role of vinculin further downstream within the complex. Vinculin is known to support the adherens junctions by stabilizing the α-catenin and F-actin interaction in a force sensitive manner ^48^. While vinculin’s mechanosensitive role within the adherens junction is acknowledged ^35,36^, its involvement in supporting cell-cell contacts under cyclic stretch, as well as its duration of action in reinforcing the adherens junction, remain less clear.

We conducted a new set of experiments to test the effect of 0.1 Hz cyclic stretch on localization of vinculin. These experiments were conducted under two different time conditions: cells were fixed immediately after 0.1 Hz cyclic stretch (0 hr) or allowed to relax for 30 minutes before fixing and staining (0.5 hr) (**Fig. 4a**). No-stretch control devices were fixed and stained following the same time intervals.

**Figure 4.**
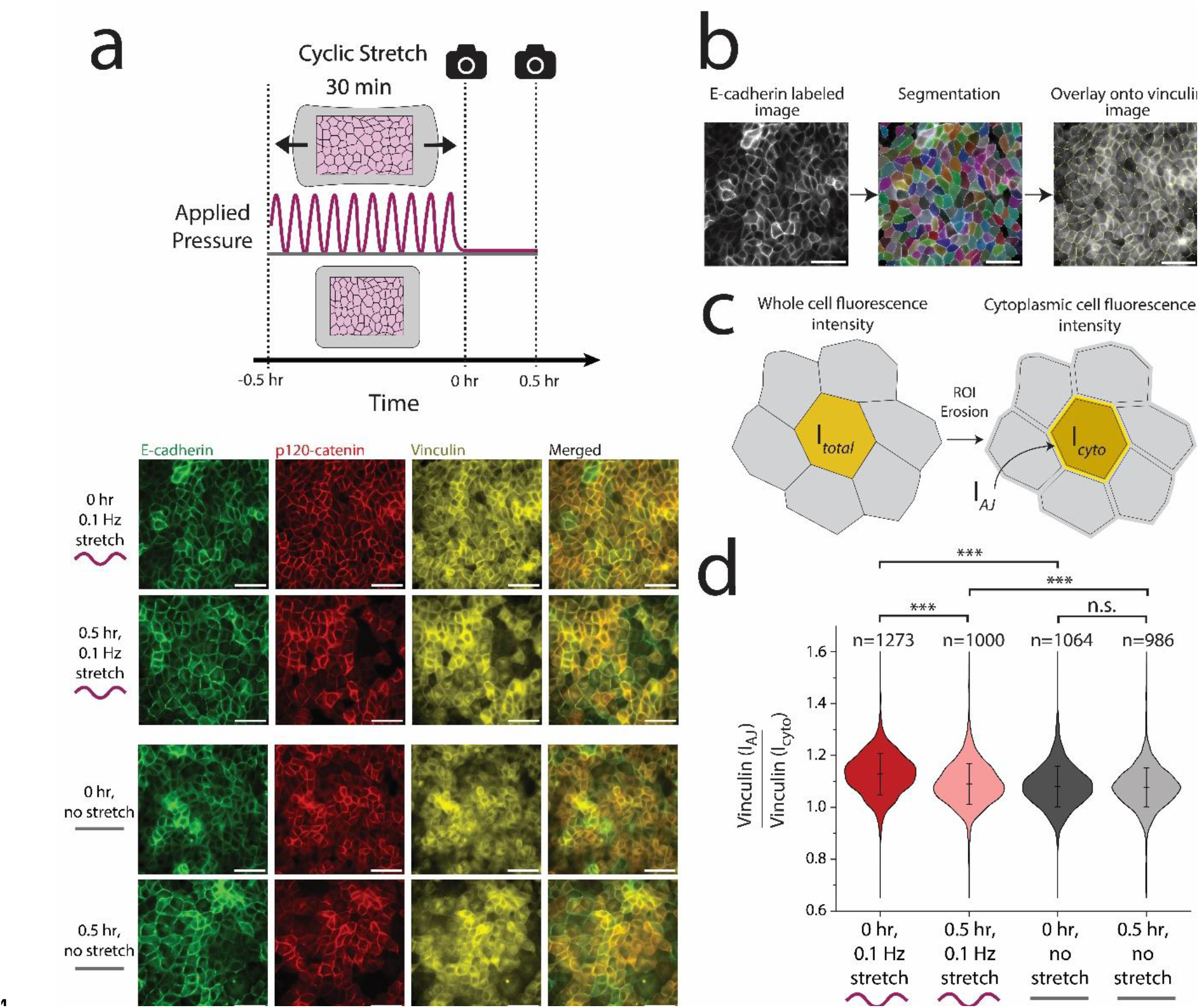
0.1 Hz cyclic stretch transiently regulates vinculin recruitment to cell-cell contacts. (a) Epithelial sheets were cyclically stretched for 30 minutes at ∼10% stretch and either fixed immediately or fixed after 30 minutes (i.e., 60 minutes after the onset of stretch). Staining for E-cadherin (green) and p120 catenin (red) showed no observable differences in response to cyclic stretch. However, staining for vinculin indicated increased localization to cell-cell contacts in response to cyclic stretch. (b) Vinculin quantification: E-cadherin labeled images were segmented and the resulting ROIs were overlaid onto the corresponding vinculin labeled images. (c) All ROIs were then eroded 3 px to isolate the cytoplasmic region of the cell from the edges of the cell. By obtaining the fluorescence intensity of vinculin in the total cell (I_total_) as well as the fluorescence intensity of vinculin in the cytoplasm for the eroded cell (I_cyto_), we computed the fluorescence intensity of vinculin at the cell-cell contact (I_AJ_) using the area fraction of the cytoplasm (see **Supplementary Fig. S5**). The ratio of vinculin intensity at cell edges to vinculin intensity in the cytoplasm showed a stark increase immediately after cyclic stretch, though diminished after 30 minutes. Scale bars are 50 µm. IHC images shown here were contrast enhanced to help visualize the proteins of interest, but not altered for segmentation analysis.

We found that while vinculin was predominantly cytoplasmic in the control condition, its localization increased at cell-cell contacts immediately after 0.1 Hz cyclic stretch. Interestingly, this intercellular vinculin recruitment was short-lived, with significant vinculin loss from cell-cell contacts after just 30 minutes. However, these levels remained higher than control levels.

### High-throughput quantification of vinculin localization

We quantified vinculin recruitment to cell-cell contacts using a cell segmentation approach. Traditional fluorescence intensity measurements for quantifying protein recruitment at cell-cell contacts are often low-throughput and subject to user bias. In contrast, cell segmentation offers increased throughput, enhanced statistical power, and reduces user bias (**Fig. 4b**). E-cadherin GFP labeled MDCK cells were segmented, and their shape outlines were then overlaid onto the corresponding vinculin-stained images of the same cells. The outlines were eroded either 0 or 3 pixels to encompass the entire cell or isolate the cytoplasmic region, respectively (**Fig. 4c**). By measuring the mean fluorescence intensity of both the entire cell and the cytoplasmic region (I_total_ and I_cyto_, respectively), then calculating the area fraction of the eroded region (**Fig. S5**), we calculated the mean fluorescence intensity of vinculin at the cell-cell contact (I_AJ_) for nearly every cell across all images.

Our results indicate that in a homeostatic state without applied mechanical stretch, a slightly higher I_AJ_/I_cyto_ ratio exists (∼ 1.08, **Fig. 4d**). Following 0.1 Hz cyclic stretch, this ratio increased to ∼1.13, signifying a robust recruitment of vinculin to cell-cell contacts (**Fig. 4d**). Notably, vinculin recruitment was transient, with a significant reduction in I_AJ_/I_cyto_ (∼1.09) after just 30 minutes of cell relaxation. However, I_AJ_/I_cyto_ vinculin levels remained statistically higher than in the control condition.

### Vinculin is necessary to suppress migration in response to cyclic stretch

With evidence of vinculin recruitment to cell-cell contacts under 0.1 Hz cyclic stretch, we tested whether vinculin was necessary for this mechanically regulated migration change. We repeated migration studies using a vinculin knock-out (KO) cell line, made in the MDCK GII background ^49^. Vinculin KO cells were observed after 30 minutes of 0.1 Hz cyclic stretch, as well as in a no-stretch control. We noted that the no-stretch control Vinculin KO cells moved slightly quicker (18-20 µm/hr) than the WT MDCK cells, consistent with other studies ^50,51^. However, Vinculin KO cells observed after 0.1 Hz uniaxial mechanical stretch slightly increased in movement, though not statistically significantly (**Fig. 5b and 5c**). This trend contrasted with the migration reduction in the MDCK WT cells after cyclic stretch. Across all timepoints, there was a tight migration band with a sigmoidal shape, indicating little variation in the speed of the cells whether subjected to mechanical force or not (**Fig. 5d-f**). These migration trends were independent of direction (**Fig. 5e, f**).

**Figure 5.**
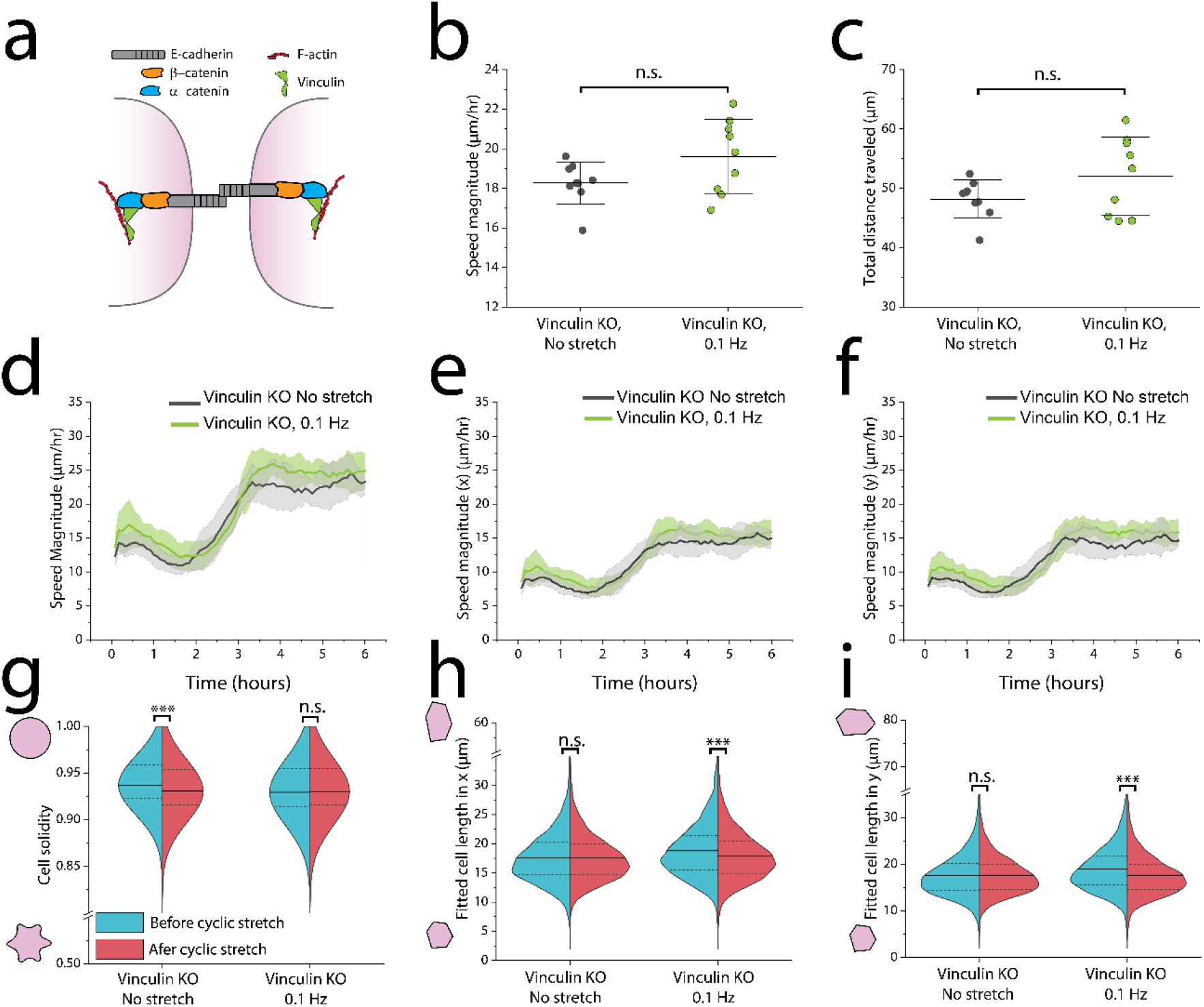
Vinculin regulates migratory response of MDCK cells to cyclic stretch, but not cell morphology. (a) Vinculin reinforces the α-catenin/F-actin complex during mechanical load. (b, c) Migration speed averaged across 6 hours following 30 minutes of cyclic stretch indicates no significant differences. (d-f) When plotted as a function of time, the Vinculin KO cells followed the same speed profile independently of cyclic stretch and direction. (g-h) Across all experiments, 5,000-10,000 cells were segmented immediately before and after 30 minutes of 0.1 Hz cyclic stretch. (g) Immediately after cyclic stretch, Vinculin KO cells retain their solidity while the solidity significantly decreases in the no-stretch control. (h, i) Vinculin KO cells also shorten their length in both x and y directions. No-stretch control: n=9 across 3 independent experiments, 0.1 Hz: n=9 across 3 independent experiments. n>5,000 cells per half of each violin plot.

IHC staining of devices after the 6-hour observation period also indicated a shift towards more cytoplasmic p120-catenin (**Fig. S6**). These results suggest that vinculin regulates recruitment or stabilization of other adapter proteins at the adherens junction.

We also labeled plasma membranes prior to cyclic stretch to observe changes in cell morphology (**Fig. 5g—i)**. Strikingly, the Vinculin KO cells retained their solidity similarly to WT MDCK cells in response to 0.1 Hz cyclic stretch (**Fig. 5g**). The Vinculin KO cells also shortened their average length in *both* the (x) and (y) directions (**Fig. 5i**).

In conclusion, these findings support vinculin’s role in regulating epithelial migration and cell length changes in response to low frequency cyclic stretch, while its role in regulating solidity was not significant.

## Discussion

After discovering that slower frequencies (0.1 Hz) regulated cell shape and migration, we investigated the adherens junction as a mechanosignaling center. Using E-cadherin T151 mutants, Vinculin KO lines, and immunohistochemistry, we confirm 3 unique findings: i) vinculin is recruited to cell-cell contacts under 0.1 Hz cyclic stretch, ii) this vinculin recruitment is transient and reduces quickly, iii) and the migratory response of the epithelium at 0.1 Hz depends on vinculin.

Our work extends prior studies on cyclic cell stretching by focusing on how different frequencies impact epithelial cell morphology and migration. While higher frequencies (1 Hz and 0.5 Hz) did not significantly alter epithelial cell migration speeds, subjecting cells to 0.1 Hz cyclic stretch for 30 minutes significantly decelerated epithelial movements over a 6-hour observation period.

By quantifying changes in cell morphology and migration, we obtained new information about collective epithelial behavior during cyclic stretch. Previous studies reported that epithelial cells reorient their shapes in response to cyclic stretch, even among different conditions ^15,17^. Our results corroborate this effect, but also demonstrate a frequency dependence.

We also found that cells retained their cell solidity at 0.1 Hz, while cell solidity decreased in the no-stretch control and higher frequency stretch conditions. Increased cell solidity is generally associated with more solid-like epithelia ^52^, making the reduced epithelial movements following 0.1 Hz stretch consistent with the retention of high cell solidity. Furthermore, we conclude that vinculin does not regulate the change in solidity. We noted that cells also experienced unique shape changes at 1 Hz that did not occur in other frequencies (**Fig. S2**). These changes included increased cell circularity and increased cell shape index, suggesting increased cell fluidity.

By conducting experiments with cells expressing a mutant E-cadherin protein (T151 cells), we demonstrated that cell migration arrest following 0.1 Hz depends on intact E-cadherin interactions at the adherens junction. Previous studies have shown that E-cadherin regulates force transfer between cells to regulate directed migration ^31,53^. We build on these studies by extending its necessity for a response to cyclic stretch via reducing cell migration.

To understand this finding, we characterized the role of vinculin in response to cyclic stretching. While vinculin plays a role in stabilizing the adherens junction, it also helps stabilize the focal adhesions of cell-ECM contacts ^54^. Recently, vinculin has been found to have an antagonistic relationship between the two regions of the cell ^35^, i.e., by perturbing vinculin expression at cell-cell contacts, vinculin at cell-ECM contacts can be disrupted. Therefore, vinculin recruitment to cell-cell contacts observed after 0.1 Hz cyclic stretch may dysregulate vinculin at cell-ECM contacts and helps arrest cell movements. This connection between protein regulation at cell-cell junctions and cell-ECM junctions is worth exploring in future studies.

An important tool we leveraged for this study was high-throughput cell segmentation. Not only did this approach significantly boost our statistical power in morphology measurements, but offered a robust method to quantify vinculin expression at cell-cell contacts with minimal user bias. For future studies (by us or others), we recommend a similar approach for quantifying expression of proteins at cell-cell contacts. The user only needs a membrane label for cell segmentation. Several computational and analytical tools (e.g., Cellpose, ImageJ, etc.) are open source.

We present here a study that i) confirms a role for different mechanical stretch frequencies in regulating collective epithelial behavior and ii) presents a novel role for vinculin in reinforcing cell-cell contacts under cyclic stretch. These results help elucidate differences observed across other cyclic stretch studies while helping understand the role of mechanical cues in regulating epithelial function.

## Materials and Methods

### Cell Culture and Cell Seeding

We used MDCK GII cells, Vinculin KO, and T151 mutant cells lacking the extracellular domain of E-cadherin both in the same MDCK GII background, as reported previously ^42,47,49^. GFP E-cadherin MDCK and Vinculin KO cells were cultured in low glucose DMEM (*ThermoFisher, 11885084*) supplemented with 10% fetal bovine serum and 1% penicillin-streptomycin at 37° C with 5% CO2. Approximately 400,000 cells in 400 µL of cell culture media were seeded in the device 36-48 h prior to experiments to create densely packed confluent monolayers.

Approximately 15 hours prior to experiments, culture media was replaced with phenol-red-free homemade basal medium (see full reagent list in SI) supplemented with 10% fetal bovine serum, 1% penicillin-streptomycin, and 50 mM HEPES to buffer the cell culture media during long-term microscopy.

T151 MDCK E-cadherin mutant cells were cultured under the same conditions, except for the addition of 20 ng/mL of doxycycline to the culture media. Addition of doxycycline represses the genetically modified promoter region of the mutant gene to maintain a wilt-type phenotype in culture conditions ^47^. For experiments, doxycycline was removed from the media during device seeding, approximately 36 hours before the experiment.

### Cyclic stretching experiments

The MDCK epithelium was imaged and cyclically stretched in a temperature-controlled chamber (37°C) of a Zeiss AxioObserver 7 inverted microscope. To cyclically stretch the epithelium, a vacuum tube connected to an electronic pressure controller (from *Red Dog Research*) was inserted into the hole in the vacuum chamber of the device. The controller is programmed to apply 0 to 60 kPa of pressure in a sinusoidal wave at variable frequencies (0.1 Hz, 0.5 Hz, or 1 Hz depending on the experiment). Immediately after cyclic stretch, the device was briefly imaged and integrated with a perfusion system for a 6-hour imaging period (see SI for details).

### High-throughput cell segmentation and ROI filtration

All images used for cell segmentation were taken using a 20x air objective (NA=0.8) and were labeled with a fluorescently tagged for E-cadherin to denote the cell-cell boundary. We utilized the CellPose “cyto” model ^55^ with a calibration of 50 pixels per image while excluding cells on edges. ImageJ’s LabelsToROIs plugin was used for shape analysis. See SI for additional post-processing details.

### Statistics

Statistics were generated from a two-tailed student t-test assuming equal variance. For PIV migration data that exhibited a non-normal distribution as determined by the Shapiro-Wilk test, we used a Mann-Whitney U test (*OriginPro 2022b, OriginLab*). P values are denoted as \**p* < .05; \*\**p* < .005; *** *p* <.0005. Dotted lines in all violin plots represent the 25^th^/75^th^ percentiles of data distribution, while the solid lines represent the mean. Shaded regions in all 6-hour migration observation plots represent the 95% CI. I-bars in all scatter plots represent the mean ± SD. For each independent experiment, we imaged 2-4 different regions across the epithelium when observing migration. For segmentation analysis, approximately 8 different regions of the epithelium were imaged.

### Antibodies

The following primary antibodies were used as previously demonstrated in MDCK cells: purified mouse anti-p120 catenin ^56^ (*BD Biosciences, 610133*) at a dilution of 1:200 and recombinant rabbit monoclonal anti-vinculin ^35^ (*abcam, ab129002*) at a dilution of 1:100. The following secondary antibodies were used, both at a 1:500 dilution: Goat anti-Mouse IgG Alexa Fluor 647 (*Invitrogen, A32728*) and Goat anti-Rabbit IgG Alexa Fluor 405 (Invitrogen, A-31556). Antibodies were diluted in PBS + 0.1% Tween20 (1X) + 1% BSA.

## Acknowledgments

The authors acknowledge funding from the NSF (1834760) and the cooperative agreement W911NF-19-2-0026 from the U.S. Army Research Office for the Institute for Collaborative Biotechnologies. LPD acknowledges the BioPACIFIC Materials Innovation Platform of the National Science Foundation (Award No. DMR-1933487) as well as useful discussions with Pruitt Lab members and M. Cristina Marchetti. The authors are also thankful to So Yomada and Sanjeevi Sivasankar (UC Davis) who gifted the Vinculin KO cells, as well as W. James Nelson who gifted the T151 and E-cadherin GFP MDCK cells. The authors acknowledge the use of the Microfluidics Lab within the California NanoSystems Institute, supported by the University of California, Santa Barbara and the University of California, Office of the President.

## Author Contributions

LPD and BLP conceived of study. LPD and BLP designed experiments. LPD and SS performed experiments and analyzed data. KM and RK contributed analytical tools for cell segmentation and PIV, respectively. LPD and SS wrote the original manuscript draft, which was reviewed and edited by LPD, SS, and BLP. BLP provided project funding and supervision.

