## Supplemental Information for "Cyclic stretch regulates epithelial cell migration in a frequency dependent manner via vinculin recruitment to cell-cell contacts"

*Device construction*

We implemented and constructed a uniaxial cell stretching device as previously reported [1]. The device consisted of two PDMS components: a thin PDMS membrane and a thicker PDMS top containing the vacuum and cell channels. The mold for the top portion was 3-D printed from RGD450 resin on a Stratasys Objet30. We then mixed PDMS (*Dow, Sylgard 184)* at a 10:1 (base: curing agent) ratio, poured it on the molds, and desiccated the PDMS for 1 hour to remove air bubbles. The PDMS was then cured on the mold at 75˚C for 4 hours. To assemble the device, we cut the cured PDMS from the mold and made 3 holes using a 1 mm biopsy punch. One hole was punched in each end of the middle chamber for inserting cells and cell media. A third hole was punched into one side of the outer chamber for the vacuum. The thin PDMS membrane (~125 µm/.005”, *Specialty Manufacturing Inc.)* was cut to the dimensions of the device top (1” x 3”), washed with 70% ethanol, then air dried with $N_2$. Using a handheld plasma wand (*Electro-Technic Products*, *BD-20AC*), we plasma treated the PDMS membrane for 1 minute, the PDMS top for 30 seconds, then the membrane for another 30 seconds. Immediately after plasma treatment, the top of the device was placed on the membrane to seal the channels. We applied 5 minutes of light pressure between the two PDMS bodies and reinforced the bond by placing the complete device in a 75˚C oven for 5 minutes. We then incubated the device with 50 µg/mL of collagen I (*Corning, 354236*) into the middle chamber overnight at 22˚C. The collagen was rinsed out with PBS before storing at 4˚C until cell seeding (approximately 2-24 hours later). Prior to cell seeding, devices were UV sterilized for 10 minutes in the biosafety cabinet.

*Cyclic stretching experiments and migration*

The vacuum was supplied by a standard in-house vacuum line. Following 30 minutes of stretch, we vented the vacuum line to remove excess pressure and removed the vacuum tube prior to imaging. Approximately 8 images (488 nm and phase contrast) of the monolayer were taken before and after cyclic stretch with both 10x and 20x objectives. We utilized a custom 3-D printed holder to prevent the device from sagging during imaging, which also kept the membrane suspended during cyclic stretch. After post-stretch imaging, we immediately set up the perfusion system for our 6-hour observation period. First, we placed the device on a 500 µm thick glass slide (*SPI Supplies, 01018T-AB)* to keep the membrane from drifting out of plane during perfusion. A syringe pump perfused cell media (as described in *cell culture and cell seeding)* through 1/32” inner diameter Tygon tubing (*McMaster-Carr, 6546T23*) into and out of the main cell channel before traveling into a waste beaker. 90-degree blunt needles *(McMaster-Carr, 75165A65)* connected the tubing to the channel holes via Luer Lock connections. The syringe pump was set to a rate of 0.3 mL/h. At the onset of perfusion, we began imaging the device. Each device was imaged at 3 regions of the channel every 5 minutes for 6 hours with a phase contrast 10x objective.

*Particle Image Velocimetry analysis*

The resulting phase contrast images from the 6-hour timelapse were aligned to remove thermal drift during imaging (*ImageJ*, linear stack alignment with SIFT). The images were then cropped to 1700 by 1150 pixels before being analyzed with PIV (MATLAB *PIVlab 2.58,* The MathWorks). PIVlab used particle image velocimetry to determine the movement of the cells within the images. The analysis in PIVlab used a FFT Window Deformation algorithm with a Gauss 2x3 point sub-pixel estimator. The interrogation pass sizes were 200, 100, and 50 for each of the three passes respectively, and each had a 50% overlap per step. In post processing vector validation, we used a standard deviation filter of 4 and a local median filter of 5. The velocity data was averaged over each frame using original MATLAB code. Migration distances were calculated using averaged velocities and the time between images.

*Immunohistochemistry and imaging of stained devices*

Devices were fixed after 4 different conditions: i) Immediately after 30 minutes of 0.1 Hz cyclic stretch, ii) immediately after 30 minutes of no cyclic stretch, iii) 30 minutes after 30 minutes of 0.1 Hz cyclic stretch, iv) or 30 minutes after 30 minutes of no cyclic stretch. 2 devices were stained for each condition (8 devices total). For the conditions involving an extra 30 minutes of relaxation, the devices were placed in the cell culture incubator before fixing and staining. All devices were washed with PBS before being fixed for 15 minutes in 4% formaldehyde (*ThermoFisher 28908*), diluted in PBS. The formaldehyde was then thoroughly washed out with PBS. For permeabilization, we used a buffer consisting of 0.1% Triton X-100 in PBS in the devices for 5 minutes at room temperature. Permeabilization was followed by a 1 hour blocking step at room temperature, using a buffer consisting of 0.3% Tween20 (1X) and 2% BSA in PBS. Primary antibodies were incubated in the devices overnight at 4˚C, following by a wash with 0.1% Tween20 (1X) and 1% BSA in PBS. Secondary antibodies were then incubated with the devices for 1 hour at room temperature in the dark. Finally, the devices were washed with 0.1% Tween20 (1X) and 1% BSA in PBS before being stored in PBS for imaging.

All devices were imaged using a 20x air objective (NA=0.8) and a 63x oil immersion objective (NA=1.2) on a Zeiss AxioObserver 7 widefield microscope. 3 separate regions were imaged per device using the following excitation filters (Vinculin:405 nm, E-cadherin: 488 nm, Vinculin: 647 nm). Each image was aligned to the apical surface of the epithelium.

*High-throughput cell segmentation and ROI filtration*

After segmentation, the masks were saved as PNG files and run through the LablesToROIs plugin in ImageJ, which calculated the shape descriptors for each cell per image. Cells having an area below 2 $\mu m^2$ or above 2,000 $\mu m^2$ were filtered out of the data sets using a custom Python script. Per shape descriptor (e.g., area, aspect ratio, solidity) all cells were then averaged per image (~2,000 cells/image). The fitted length for cell shape measurements was determined using ImageJ's bounding rectangle feature

For Vinculin KO experiments, cell segmentation was determined using a CellMask Deep Red plasma membrane stain (*ThermoFisher, C10046*). CellMask was incubated at a 1:1000 dilution for 10 minutes before being washed out and replaced with imaging medium, after which the experiment was performed immediately.

*Analysis of vinculin localization at cell-cell contacts*

First, the E-cadherin labeled images from the 63x oil immersion objective were segmented using Cellpose, as described in *high-throughput cell segmentation*. Using the LabelsToROIs plugin in ImageJ, the segmented masks were overlaid onto the corresponding vinculin labeled images. Segmented masks were then processed under 2 separate conditions: i) masks were eroded 3 pixels to remove the cell-cell contact or ii) masks were eroded 0 pixels to contain the entire cell, including the cell-cell boundary. The corresponding mean fluorescence intensities of vinculin were calculated for each cell under each condition (i.e., the mean fluorescence intensity of vinculin in the entire cell and the mean fluorescence intensity of vinculin excluding the cell-cell contact).

To determine the mean fluorescence intensity of vinculin at the cell-cell contacts, we calculated the approximate area fraction of the 3-pixel eroded cell to be 0.89 (**Fig. S5**). This area fraction allowed us to calculate the mean fluorescence intensity at the cell-cell contacts.

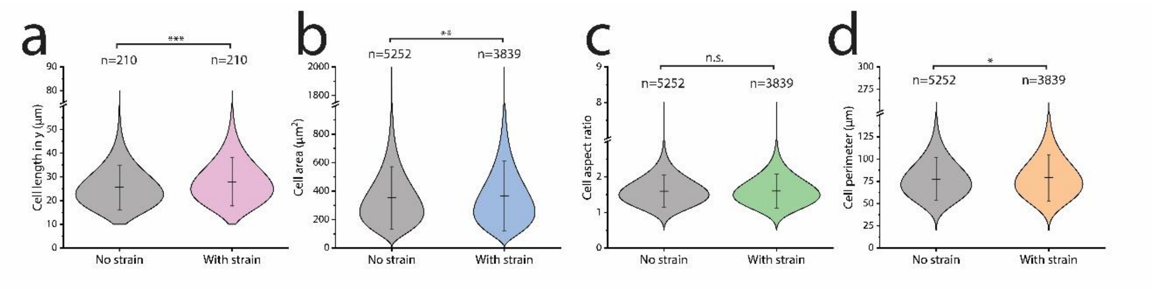


**Figure S1. Quantification of cell shape under 60 kPa static strain**. (a) For 7 separate experiments, 210 cells were manually measured along the direction of stretch ($\varepsilon_{yy}$) with and without stretch, yielding a difference in approximately 10% and compared using a paired t-test. For other cell shape descriptors (b-d), cells were segmented from 20x magnification E-cadherin GFP labeled images. Stretch significantly increased cell area and perimeter, while cell aspect ratio remained unchanged. I-bars in violin plots represent mean <u>+</u> SD. Violin plots are distributed via a gamma curve. **p* < .05; ***p* < .005; *** *p* <.0005 using a student t test.

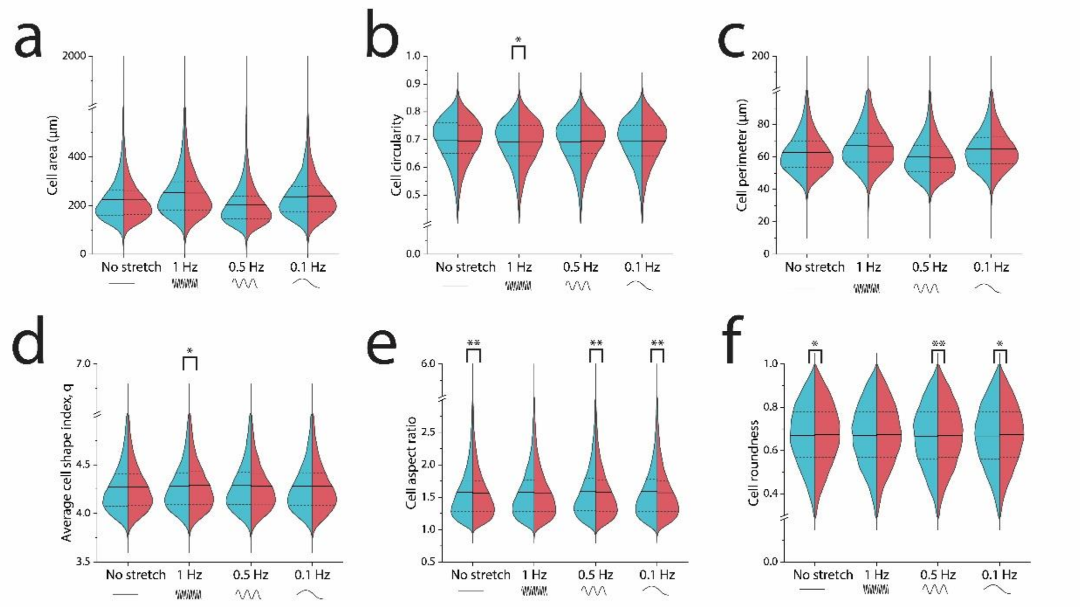


**Figure S2. Changes in cell morphology after cyclic cell stretch.** (a, c) There were no differences in average cell area nor perimeter among all cyclic stretch conditions. (b, d) However, we observed small significant changes in cell shape index, q, and cell circularity under higher frequency 1 Hz. (e, f) For aspect ratio and roundness, we observed subtle (though significant) changes in lower frequencies and the control condition. q is defined as the perimeter/sqrt(area). n>10,000 cells per half of each violin plot. Dotted lines in the violin plots represent the $25^{th}/75^{th}$ percentiles of data distribution. **p* < .05; ***p* < .005; *** *p* <.0005 using a student t test.

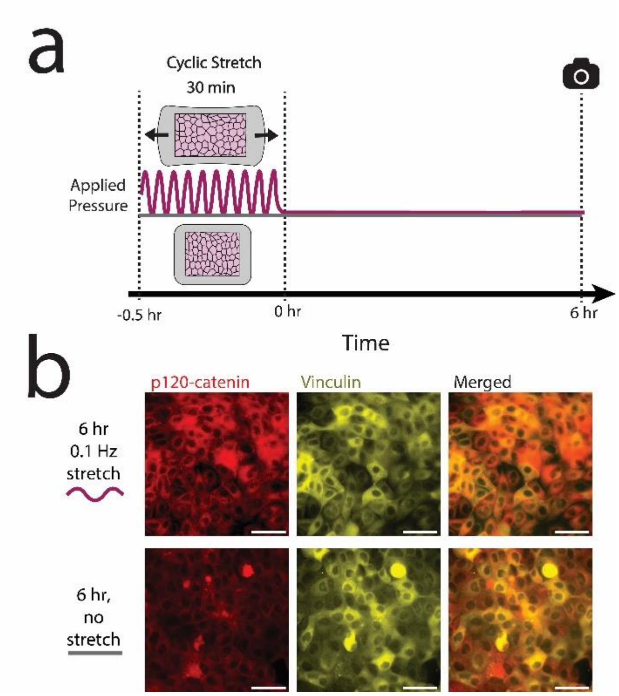


**Figure S3. p120-catenin and vinculin are predominantly cytoplasmic in T151 E-cadherin mutant cells.** No-stretched and stretched devices with the T151 cells were fixed and stained after the 6-hour observation period. There is no clear localization of p120-catenin nor vinculin at cell-cell contacts. Scale bars are 50 µm. IHC images shown have enhanced contrast to help visualize the proteins of interest.

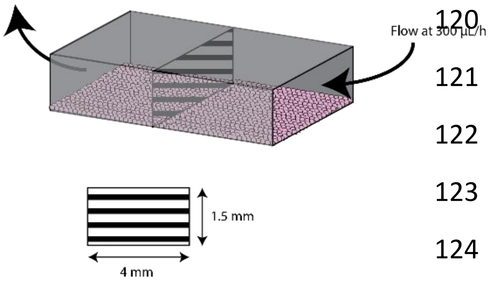


$$A = 6\,mm^2$$

$$6\,mm^2 = 6 * 10^6\,\mu m^2$$

$$Q = A * v = 300\,\frac{\mu L}{h}$$

$$300\,\frac{\mu L}{h} = (6 * 10^6 \mu m^2) * v$$

$$v = \left(\frac{300\,\mu L}{6 * 10^6\,\mu m^2 h}\right)\left(\frac{1L}{10^6 \mu L}\right)\left(\frac{10^{-3} m^3}{1\,L}\right)\left(\frac{10^6 \mu m}{1\,m}\right)\left(\frac{1\,h}{3600\,s}\right)$$

$$v = 13.89\,\frac{\mu m}{s}\,Pa = \frac{kg}{m * s^2} = P = \frac{1}{2} v^2 \rho$$

$$\rho = 1.0\,\frac{g}{mL} = \frac{1.0\,g}{10^{-12} \mu m^3}$$

$$P = \frac{(13.89\,\mu m)^2 * 1.0 g}{2 * 10^{-12}\,\mu m^3 * s^2}$$

$$= 9.65 * 10^{-11}\,\frac{g}{\mu m * s^2}$$

$$9.65 * 10^{-11}\,\frac{g}{\mu m * s^2}\left(\frac{1\,kg}{1000\,g}\right)\left(\frac{10^6 \mu m}{1\,m}\right)$$

$$= 9.65 * 10^{-8} Pa$$

$$9.65 * 10^{-8} Pa\left(\frac{10\,dyne}{1\,Pa * cm^2}\right) = 9.65 * 10^{-7}\,\frac{\text{dyne}}{cm^2}$$

**Figure S4. Calculation of shear flow on epithelial monolayer within perfusion system.**

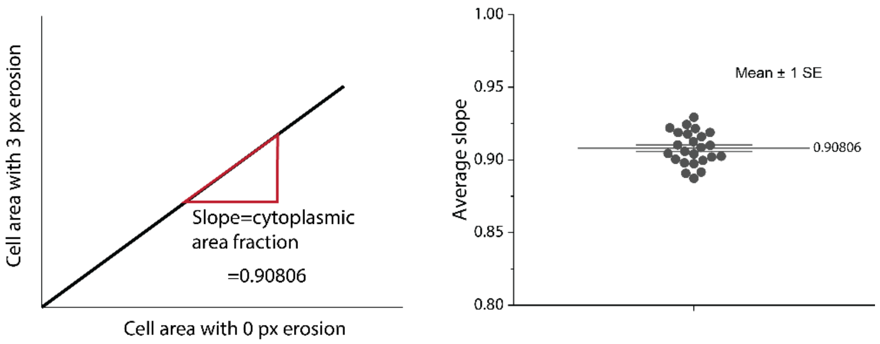


$$A_{total} = Total\ cell\ area$$
$$A_{cyto} = Total\ cytoplasmic\ area$$
$$A_{AJ} = Total\ cell\ contact\ area$$

$$A_{total} = A_{cyto} + A_{AJ}$$

$$A_{cyto} = 0.90806(A_{total})$$

$$A_{AJ} = (1 - 0.90806)(A_{total})$$

$$I = mean\ fluorescence\ intensity$$

$$I_{total} = I_{AJ}(1 - 0.90806) + I_{cyto}(0.90806)$$

$$I_{AJ} = \frac{I_{total} - I_{cyto}(0.90806)}{(1 - 0.90806)}$$

**Figure S5. Calculation of vinculin fluorescence intensity at cell-cell contacts.** (left) For each data set, the cell areas of a 0 px eroded cell were plotted against the areas for the corresponding 3 ox eroded cell. (right) The average slope of these plots was 0.90806, indicating the cytoplasmic area fraction.

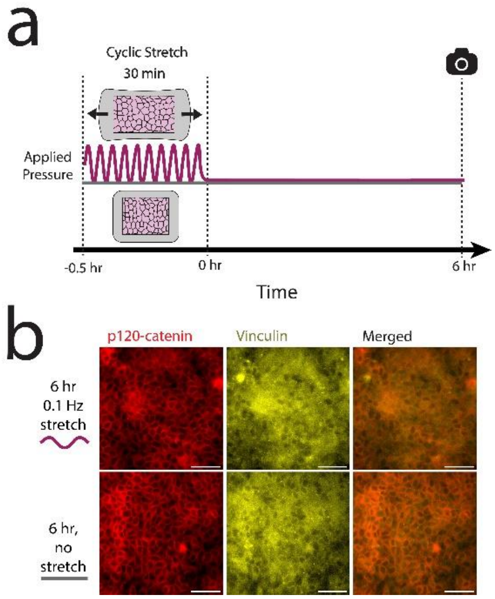


**Figure S6. P120-catenin delocalizes from cell-cell contacts in Vinculin KO cells.** No-stretch control and stretched devices with the Vinculin KO cells were fixed and stained after the 6-hour observation period. Qualitatively, p120-catenin is less localized to cell-cell contacts than for the WT MDCK cells. As expected, vinculin expression is very low (observed signal is mostly background noise). Scale bars are 50 µm. IHC images shown have enhanced contrast to help visualize the proteins of interest.

### Homemade basal medium for cell imaging experiments

The following reagents were mixed in water, pH adjusted to 7.0 using HCL, then filter sterilized with a 0.1 µm filter. Methionine, Calcium Chloride, and the MEM Vitamin Solution (100x) were added after pH adjustment.

| **Ingredient** | **Concentration (mg/L)** |
|---|---|
| KCL | 400 |
| $MgSO_4.7H_2O$ | 200 |
| NaCl | 5963 |
| D-Glucose (dextrose,monohydrate) | 1000 |
| L-arginine. HCL | 126 |
| L-cystine. 2HCL | 31 |
| L-glutamine | 296 |
| L histidine HCL. $2H_2O$ | 42 |
| L-isoleucine | 52 |
| L-leucine | 52 |
| L-lysine HCL | 73 |
| L-phenylalanine | 32 |
| L-threonine | 48 |
| L-tryptophan | 10 |

| | |
|---|---|
| L-tyrosine | 52 |
| L-valine | 46 |
| $NaHCO_3$ | 1000 |
| Na Hepes | 2603 |
| $NaH_2PO_4 . H_2O$ | 140 |
| Methionine | 167 |
| Calcium Chloride ($CaCl_2$) | 200 |
| MEM Vitamin Solution (100×) (*Sigma, M6895*) | N/A |
